# TMAO miscompartmentalization is a reversible driver of autism pathophysiology

**DOI:** 10.1101/2024.05.30.596635

**Authors:** Jean-Marie Launay, Nicolas Vodovar

## Abstract

Autism spectrum disorder (ASD) is a complex and heterogeneous neurodevelopmental disorder. Contrary to what has been reported for genetics and gut dysbiosis, ASD appears to be very homogeneous when considering tryptophan metabolism. Indeed, multiple biochemical anomalies have been observed in most individuals with ASD. Following up on these findings, we found that ASD is strongly associated with the miscompartmentalization of the chemical chaperone trimethylamine *N*-oxide (TMAO). Intracellular TMAO was markedly reduced in individuals with ASD as a result of altered fluid/electrolyte homeostasis and was responsible for numerous biochemical anomalies described in ASD. Administration of urea in a rat model of ASD that recapitulates the biochemical anomalies observed in humans not only restored biochemical parameters but also broadly improved all behaviours. Our results demonstrate the major role of TMAO in the pathophysiology of ASD and cellular physiology, although TMAO miscompartmentalization is not causal for ASD. We anticipate that urea, which is already clinically approved, offers a breakthrough therapeutic opportunity for ASD.

## Introduction

ASD is a complex and heterogeneous neurodevelopmental disorder that is characterized by impaired social communication, restrictive interest, repetitive behaviours, and hypersensitivity^1^. To date, there is no broadly effective treatment for ASD. While genetic factors have been linked to ASD^2–4^, at most 10% of cases can be attributed to specific mutations, and >1000 genetic variants have been associated with ASD^4^. Gut dysbiosis has also been associated with the severity of mental illness^5^ and has been proposed as a driving factor in ASD, although dysbiosis appears more as a consequence than a cause of ASD^6^. From a metabolic standpoint, ASD is characterized by numerous biochemical alterations, in particular in the tryptophan pathway. These alterations include reduced enzymatic activities linked to serotonin (5-HT) catabolism by sulphoconjugation^7^, melatonin^8^ and nicotinamide adenine dinucleotide (NAD^+^) production^9^. Furthermore, multiple enzymatic activities of the kynurenine pathway are also reduced^9^. In marked contrast to genetics and microbiota findings, our recent findings suggest that ASD is metabolically very homogeneous^9^. These enzymatic anomalies appear to extend beyond the tryptophan metabolism since the activity of another sulfotransferase involved in extracellular matrix homeostasis is also reduced^7^. Overall, the coexistence of these apparently unrelated biochemical alterations suggests higher-order regulatory mechanisms.

Trimethylamine *N*-oxide (TMAO) is the oxidation product of flavin-containing monooxygenase 3 (FMO3) of gut microbiota-produced trimethylamine^10^. TMAO is a chemical chaperone that facilitates protein folding^11–14^ and prevents protein denaturation by urea^15^ and hydrostatic pressure in marine animals^16^. TMAO also stabilizes RNA tertiary structure *in vitro*^17^ and increases the rigidity of lipid membranes^18–21^. More recently, increased TMOA plasma levels have been associated with cardiometabolic diseases^22–24^, and a study has reported elevated plasma TMAO levels in ASD^25^. However, little is known regarding its physiological function.

Here, we show that intracellular TMAO depletion, as a result of decreased extracellular osmolarity, is responsible for most of the biochemical alterations observed in ASD. Restoration of intracellular TMAO by urea in a rat model of ASD restored metabolic homeostasis and markedly improved all ASD-like behaviours. These results posit TMAO as a major regulator of cellular physiology and urea as a potential breakthrough treatment for ASD.

## Results

### Intracellular TMAO is lower in ASD

Melatonin is produced from 5-HT by the sequential action of aralkylamine N-acetyltransferase (AANAT) and acetylserotonin O-methyltransferase (ASMT) whose activities are reduced in ASD^8^. Both enzyme activities rely on the 14-3-3 chaperone whose levels are also reduced in ASD^8^. AANAT activity^26^ was more sensitive than ASMT activity to 14-3-3 levels^8^, which is reflected by the strong positive correlation between platelet 14-3-3 concentration and AANAT activity (***Fig. 1a***), and the lack thereof for ASMT activity (***Fig. 1b***). However, a decrease in ASMT activity without any decrease in AANAT activity was observed in ∼41% of individuals with ASD^8^, suggesting another chaperone could be involved in melatonin synthesis. Hence, we immunoprecipitated AANAT or ASMT from 50×10^9^ platelets of healthy subjects (n = 6) and subjected the immunoprecipitates to native mass spectrometry^27^. Regardless of the antigen targeted, the immunoprecipitates always contained AANAT, 14-3-3, and ASMT, indicating that the three proteins formed a complex *in vivo*. We further subjected the immunoprecipitates to GC-MS and consistently found three peaks (m/z = 75, 58, and 44), which were identified as TMAO and its fragmentation products (***Fig. 1c***).

**Fig. 1:**
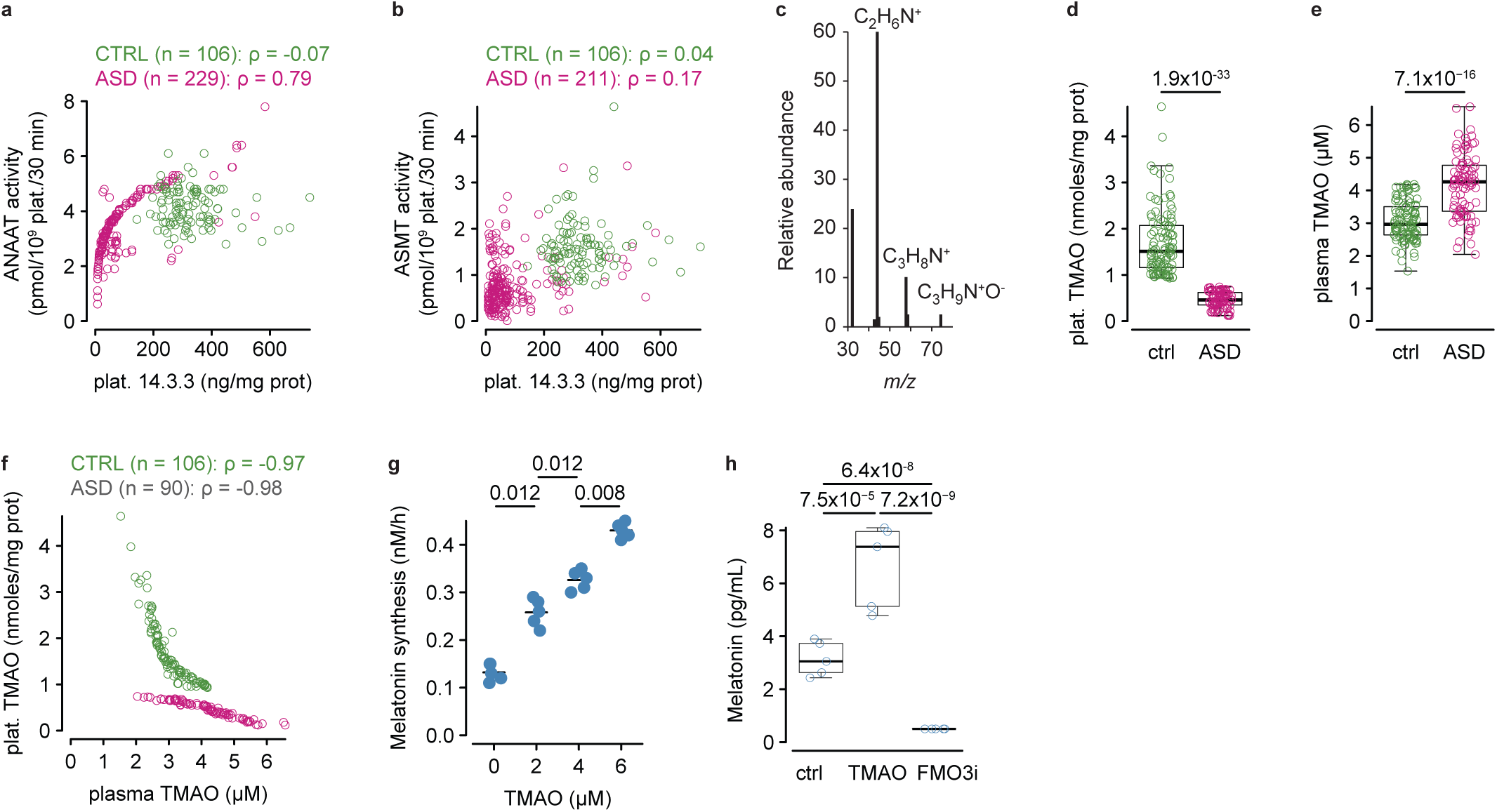
TMAO chaperones melatonin synthesis. **a,** relationship between platelet 14-3-3 concentration and platelet AANAT activity measured in the platelets of controls and individuals with ASD. **b,** relationship between platelet 14-3-3 concentration and platelet ASMT activity measured in the platelets of controls and individuals with ASD. **c,** representative GC/MS spectrum obtained from platelet AANAT or ASMT immunoprecipitates of healthy individuals. **d,** platelet TMAO concentration in controls (n = 106) and individuals with ASD (n = 90). **e,** plasma TMAO concentration in controls (n = 106) and individuals with ASD (n = 90). **f,** relationship between plasma and platelet TMAO concentrations in controls and individuals with ASD. **g,** *in vitro* melatonin synthesis by recombinant human AANAT, ASMT and 14-3-3 with respect to TMAO concentration (n = 5 per TMAO concentration). **h,** melatonin plasma levels in control mice, that successively received TMAO and FMO3 inhibitor (n = 5, see methods). Intergroup comparisons were performed using the Wilcoxon sum-rank test (d and e) or the Kruskal-Wallis test followed by the Wilcoxon sum-rank test corrected for multiple comparisons (g). Within-group comparisons were performed using repeated-measure ANOVA on log-transformed data followed by the Tukey HSD test (h). Correlations were calculated using Spearman’s correlation coefficient (ρ). The number of individuals in the cohort is indicated on each plot.

Next, we quantified the concentration of TMAO in platelets and plasma of individuals with ASD and controls from the same cohort in which we described the various anomalies in the tryptophan metabolism^7,9,28^. Platelet TMAO was markedly lower (***Fig. 1d***), while plasma TMAO was higher (***Fig. 1e***) in individuals with ASD with strong correlations between plasma and platelet TMAO in both ASD and controls (***Fig. 1f***), indicating TMAO miscompartmentalization in ASD. To test the role of TMAO in melatonin synthesis, we measured melatonin production *in vitro* by incubating 5-HT with human recombinant AANAT, ASMT, 14-3-3 and TMAO, and found that melatonin synthesis increased with increasing TMAO concentrations (***Fig. 1g***). In male mice (FVB/N), the administration of TMAO (50 mg/kg) provoked an increase in platelet and plasma TMAO (***Extended Fig. 1***), and plasma melatonin (***Fig. 1h***), while the administration of an FMO3 inhibitor (3mg/kg phenylthiourea) in the same mice after a one-week wash-out period, decreased platelet and plasma TMAO (***Extended Fig. 1***), and plasma melatonin (***Fig. 1h***). Collectively, these data suggest that TMAO miscompartmentalization in ASD is responsible for a decrease in ANAAT/ASMT/14.3.3 chaperoning by TMAO, hence reduced melatonin production.

### TMAO and biochemical alterations in ASD

The eukaryotic tryptophan metabolic pathways (***Fig. 2a***) are markedly altered in ASD. Besides deficits in melatonin synthesis^8^ and sulphoconjugation (PST-M and PST-P)^7^, several alterations of the kynurenine pathway have been observed, namely a decrease in 3-hydroxyanthranilate oxidase (3-HAO) activity which resulted in 3-hydroxyanthranilate (3-HAA) accumulation, and a deficit in NAD^+^ production^9^. As TMAO is a chemical chaperone, and based on the principle of parsimony, we tested the role of TMAO on those alterations *in vitro* using purified enzymes, and *in vivo* using the exogenous TMAO/FMO3i mouse model. First, PST-P activity increased with increasing TMAO concentrations added to the reaction buffer (***Fig. 2b***) and PSTs – the sole PST in rodents, activity was unaffected by TMAO administration but decreased when TMAO synthesis was blocked by FMO3i *in vivo* (***Fig. 2c***). Second, 3-HAO activity, estimated by the plasma quinolinic (QA) + picolinic acids (PA)/3-HAA ratio (***Fig. 2a***), was strongly negatively correlated with plasma TMAO, only in individuals with ASD (***Fig. 2d***). *In vitro*, the activity of human recombinant 3-HAO increased with increasing TMAO concentrations added to the reaction buffer (***Fig. 2e***), and *in vivo* exogenous TMAO increased while FMO3i decreased liver 3-HAO activity (***Fig. 2f***). Finally, plasma NAD^+^ concentrations negatively correlated with plasma TMAO, only in individuals with ASD (***Fig. 2g***). In mice, NAD^+^ plasma levels increased after TMAO administration and decreased after FMO3 inhibition (***Fig. 2h***). Taken together, these results indicate that TMAO chaperones numerous enzymatic processes that were found altered ASD^7–9^.

**Fig. 2:**
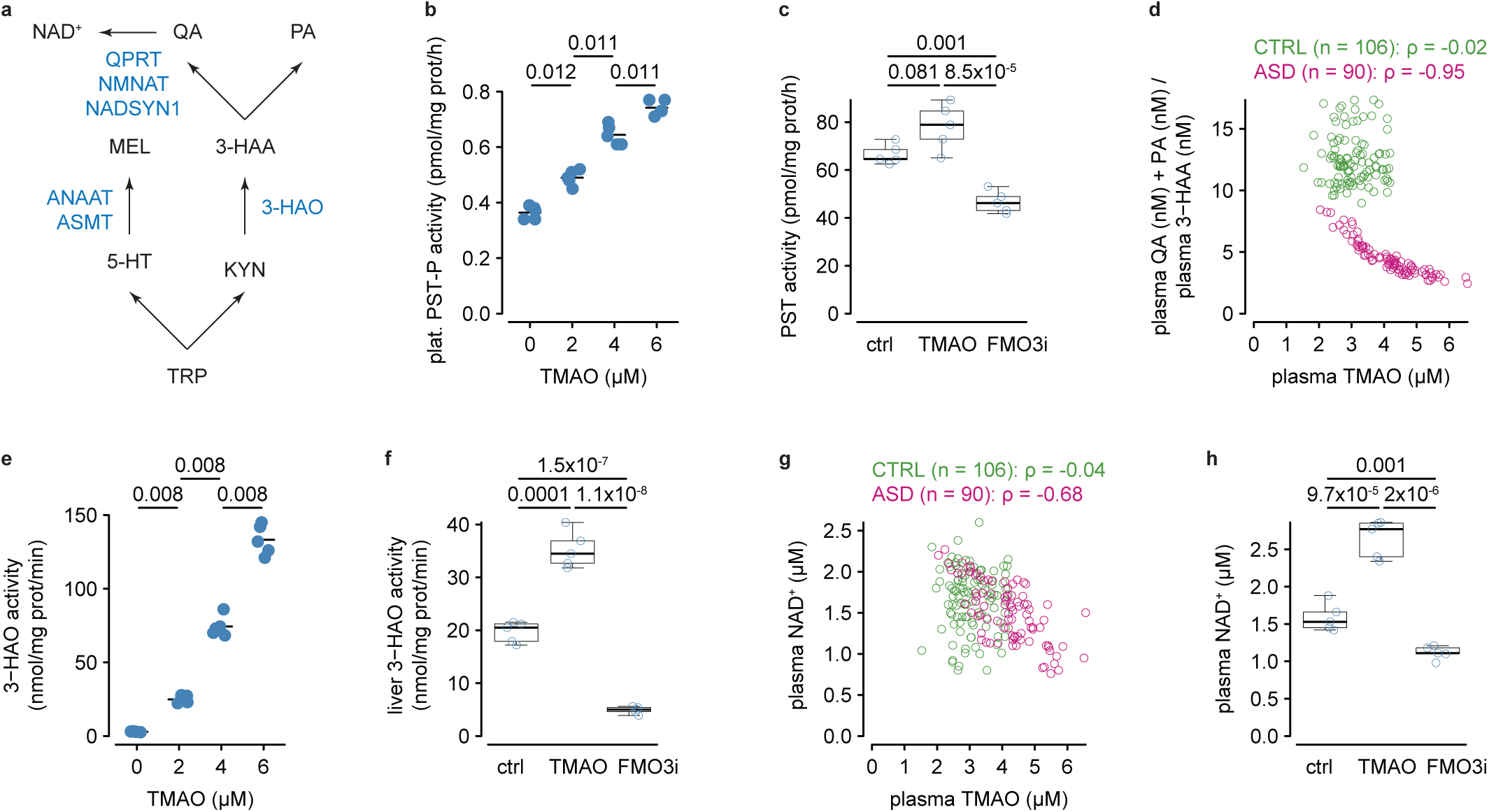
TMAO chaperones numerous enzymatic activities. **a,** schematic representation of tryptophan (TRP) metabolism relevant for the study. MEL: melatonin; KYN: kynurenine, 3-HAA: 3-hydroxyanthranilic acid, QA: quinolinic acid, PA: picolinic acid, NAD^+^: nicotinamide dinucleotide. Enzymes are indicated in blue: QPRT: quinolinate phosphoribosyl transferase, NMNAT: nicotinamide nucleotide adenylyl transferase, NADSYN1: glutamine-dependent NAD^+^ synthetase. **b,** activity of recombinant human phenol-specific phenol sulphotransferase (PST-P) with respect to TMAO concentration (n = 5 per TMAO concentration). **c,** platelet PST activity measured in control mice, that successively received TMAO and FMO3 inhibitor (n = 5, see methods). **d,** relationship between plasma TMAO concentration and 3-hydroxyanthranilate oxidase (3-HAO) activity estimated by the QA+PA/3-HAA ratio in controls and individuals with ASD. **e,** activity of recombinant human 3-HAO with respect to TMAO concentration in the reaction buffer (n = 5 per TMAO concentration). **f,** liver 3-HAO activity measured in control mice, that successively received TMAO and FMO3 inhibitor (n = 5, see methods). **g,** relationship between plasma TMAO and NAD^+^ in controls and individuals with ASD. **h,** plasma NAD^+^ measured in control mice, that successively received TMAO and FMO3 inhibitor (n = 5, see methods). Intergroup comparisons were performed using the Kruskal-Wallis test followed by the Wilcoxon sum-rank test corrected for multiple comparisons (b and e). Within-group comparisons were performed using repeated-measure ANOVA on log-transformed data followed by the Tukey HSD test (c, f, and h). Correlations were calculated using Spearman’s correlation coefficient (ρ). The number of individuals in the cohort is indicated on each plot.

### Osmolarity and intracellular depletion of TMAO

In sharks, TMAO can be released as a compatible solute from cells to compensate for a decrease in extracellular osmolarity^29,30^. In our cohort, the only osmolarity-related parameter available was natremia, and we found that 81% (n = 73/90) of individuals with ASD were hyponatremic (< 135 mM, ***Fig. 3a***). Furthermore, there were strong correlations between natremia and platelet (***Fig. 3b***) or plasma (***Fig. 3c***) TMAO, only in individuals with ASD and especially in hyponatremic individuals. To confirm the role of osmolarity in TMAO compartmentalisation, we resuspended fresh human platelets in Tyrode solutions at different osmolarities by dilution (300 mOsm/L and 285 mOsm/L are within the normal range) and measured TMAO in platelets and the supernatant. At 300 mOsm/L, there was no detectable TMAO release, while it was released at lower osmolarity in a dose-response fashion (***Fig. 3d***). To confirm the impact of osmolarity-driven TMAO cellular release, we measured platelet ANNAT and ASMT activities in the same platelet samples and found that both activities decreased with decreasing osmolarities (***Fig. 3e***).

**Fig. 3:**
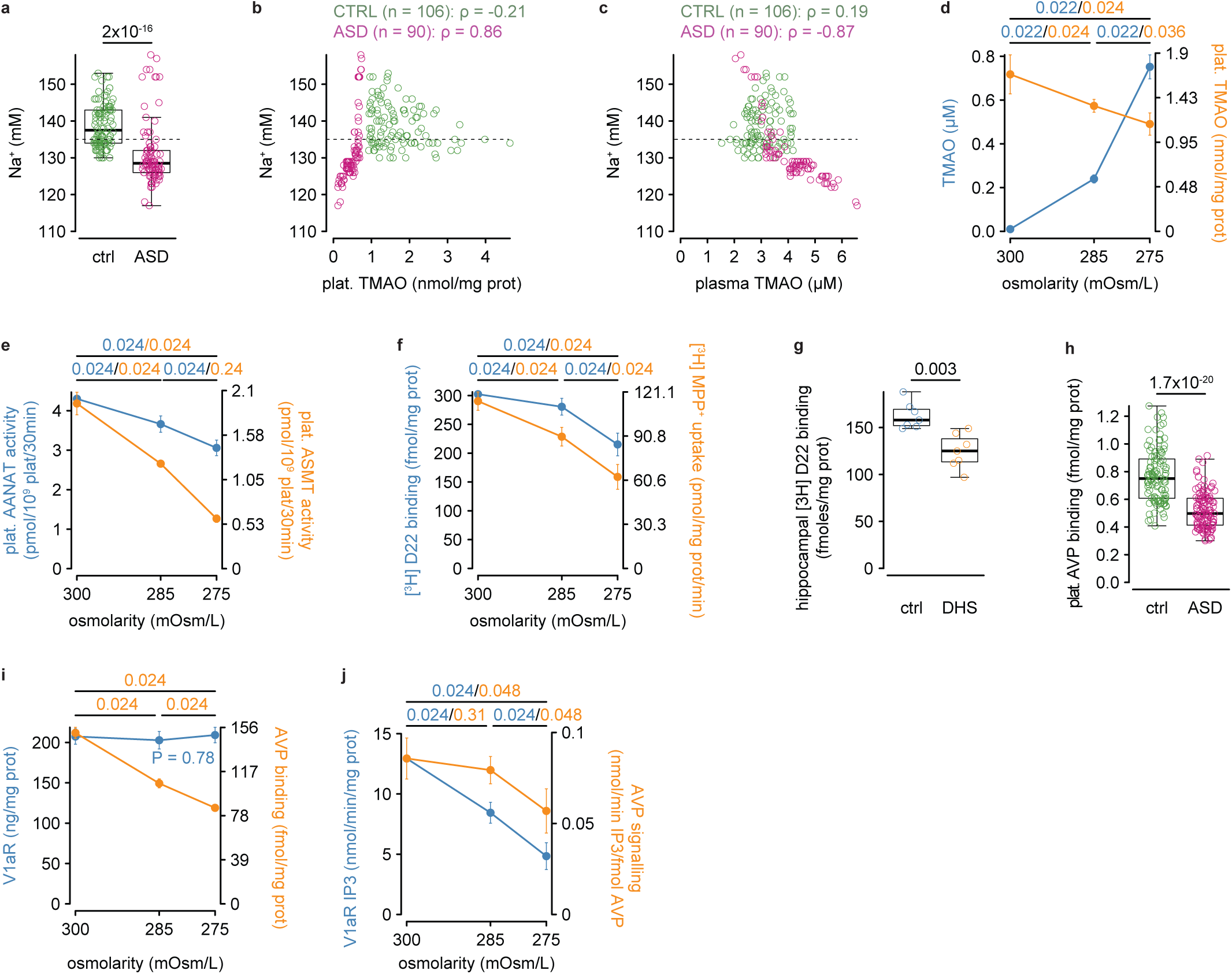
Osmolarity drives TMAO efflux. **a,** natremia of controls (n = 106) and individuals with ASD (n = 90). **b,** relationship between plasma TMAO and natremia in controls and individuals with ASD. **c,** relationship between platelet TMAO and natremia in controls and individuals with ASD. **d,** Tyrode (blue) and platelet (orange) TMAO concentrations measured in human platelets incubated in Tyrode solutions of varying osmolarity (n = 5 per condition). **e,** platelet ANAAT (blue) and ASMT (orange) activities measured in human platelets incubated in Tyrode solutions of varying osmolarity (n = 5 per condition). **f,** [^3^H] D22 binding (blue) and [^3^H] MPP+ uptake (orange) measured in human platelets incubated in Tyrode solutions of varying osmolarity (n = 5 per condition). **g,** [^3^H] D22 binding on the hippocampus of control and DHS rats (n = 7 per condition). **h,** AVP binding on platelets of controls (n = 106) and individuals with ASD (n = 90). **i,** platelet V1aR concentrations (blue) and AVP bindings (orange) measured in human platelets incubated in Tyrode solutions of varying osmolarity (n = 5 per condition). **j,** platelet IP_3_ produced by V1aR after incubation with AVP (blue) and relative to AVP binding (orange) activities measured in human platelets incubated in Tyrode solutions of varying osmolarity (n = 5 per condition). Intergroup comparisons were performed using the Wilcoxon sum-rank test (a and h) or the Kruskal-Wallis test followed by the Wilcoxon sum-rank test corrected for multiple comparisons (d-f and i-j). Correlations were calculated using Spearman’s correlation coefficient (ρ). The number of individuals in the cohort is indicated on each plot. In a-c, the dotted line indicates the lower limit of the reference range.

While TMAO can be released by cells via various transporters, its cellular uptake is predominantly performed by the organic cation transporter 2 (OCT-2)^31^. While TMAO release is increased, its cellular uptake could also be affected. We therefore incubated human platelets in Tyrode solutions at different osmolarities with the pan-OCT inhibitor, decynium-22 (D22), and found a decrease in D22 binding with decreasing osmolarity (***Fig. 3f***). *In vivo*, we also found a decrease in D22 binding in the hippocampus of DHS rats, a model of ASD^32^ (***Fig. 3g***). To functionally test a deficit in OCT-2 transport, we measured the transport of 1-methyl-4-phenyl pyridinium (MPP^+^) in human platelets resuspended in Tyrode solutions at different osmolarities and found a decrease in MPP^+^ uptake (***Fig. 3f***). Importantly, the binding of MPP^+^ to OCT-2 is dependent on cholesterol^33^, and TMAO has membrane rigidifying properties similar to cholesterol, yet by a different mechanism^18^. Hence, we further investigated the binding of arginine-vasopressin (AVP) to its receptor, as the high affinity of the oxytocin-AVP receptor family for their ligands is dictated by cholesterol^34^. Furthermore, a recent small clinical trial found that intranasal AVP administration to children with ASD was more beneficial to children with the highest pre-treatment AVP blood levels, suggesting that AVP receptors could be less functional^35^. First, we measured AVP binding to V1aR – the sole AVP receptor present in platelets^36^, in platelet samples from our cohort and found a lower AVP binding in ASD (***Fig. 3h***) despite similar V1aR protein levels compared to controls (***Extended Fig. 2***). Furthermore, AVP binding but not V1aR levels decreased with osmolarity in platelet from healthy subject incubated in Tyrode solutions of varying osmolarities (***Fig. 3i***). This decrease in AVP binding was associated with an overall decrease in IP_3_ production, even when IP_3_ production was expressed relatively to AVP binding, when osmolarity reached 275 mOsm/L (***Fig. 3j***). Taken together, these results strongly suggest that TMAO miscompartmentalization in ASD results from altered fluid/electrolyte homeostasis and has a broad impact on cell physiology since transport as well as GPCR binding and signalling also appear to be influenced by TMAO.

### Restoration of osmolarity/natraemia by urea in young DHS rats

We next hypothesised that restoring fluid/electrolyte balance in ASD using a compatible solute should correct TMAO compartmentalisation and alleviate the associated biochemical burden. To this end, we used the rat developmental hyperserotonaemia (DHS) model of ASD^32^ (***Fig. 4a***) and urea as a compatible solute, as it is clinically approved for the treatment of acute or chronic hyponatraemia^37,38^. Importantly, DHS rats shared numerous biochemical alterations found in individuals with ASD from the cohort: *i)* they were moderately hyponatremic (< 130mM, ***Fig. 4b***), *ii)* had lower platelet and higher plasma TMAO levels (***Fig. 4c-d***), *iii)* exhibited hyperserotonemia (***Fig. 4e***) as previously described^32^, and *iv)* lower plasma levels of melatonin (***Fig. 4f***) and NAD^+^ (***Fig. 4g***). Young DHS rats were administered 0.5g/kg/day urea subcutaneously for five days from PDN21 to PDN25 using an osmotic pump (***Fig. 4a***). At PDN33, urea had restored TMAO compartmentalisation and all the biochemical parameters tested to control levels (***Fig. 4b-g***).

**Fig. 4:**
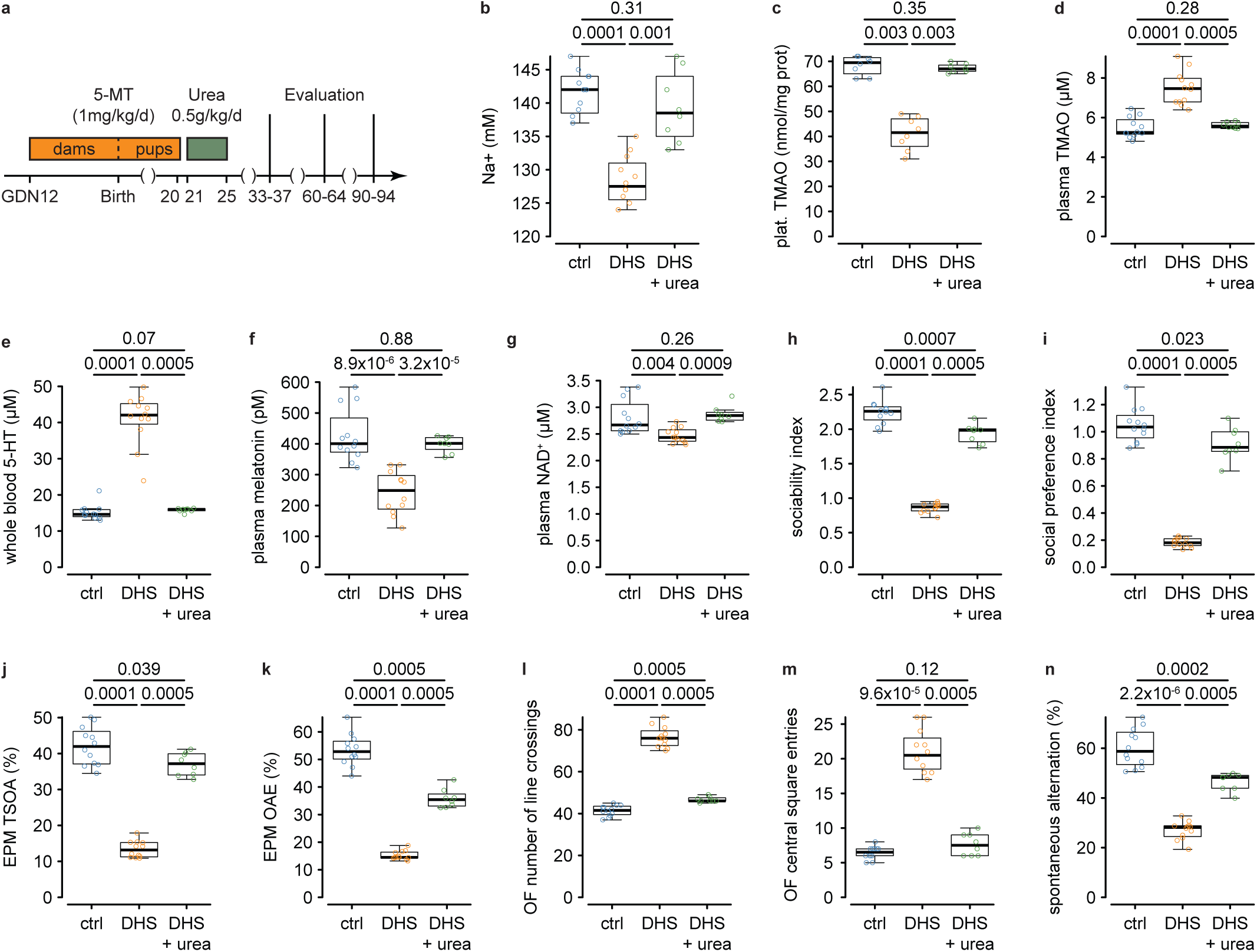
Effects of urea on young DHS rats. **a,** schematic representation of the experimental protocol. Natremia (**b**), platelet (**c**) and plasma (**d**) TMAO, whole-blood serotonin (5-HT, **e**), plasma melatonin (**f**) and NAD^+^ (**g**) concentrations in control rats (n = 12, blue), DHS rats (n = 12, orange) and DHS rats treated with urea at PDN21-PDN25 (n = 8, green). Sociability index (**h**), social preference index (**i**), time spent in the open arm (TSOA, **j**) and the number of open arm entries (OAE, **k**) in the elevated plus maze (EPM), the number of line crossing (**l**) and the number of central square entry (**m**) in the open field (OF), and the % of spontaneous alternation (**n**) in control rats (n = 12, blue), DHS rats (n = 12, orange) and DHS rats treated with urea (n = 8, green). Intergroup comparisons were performed using the Kruskal-Wallis test followed by the Wilcoxon sum-rank test corrected for multiple comparisons. 5-MT: 5-methoxytryptamine.

From a behavioural standpoint, at PND33, DHS rats exhibited a decrease in sociability using the three-chamber test for sociability and social preference (***Fig. 4h-i***), an increase in anxiety using the elevated plus maze (***Fig. 4j-k***), increased locomotion in the open field tests (***Fig. 4l-m***), and an increase in repetitive tasks using the spontaneous alternation test (***Fig. 4n***). Urea administration markedly improved all behavioural parameters tested (***Fig. 4h-n***). When the same rats were re-evaluated at PDN60-64 and PDN90-94, all biochemical but natremia and all behavioural parameters gradually reverted toward levels observed in untreated DHS rats (***Extended Fig. 3***). Altogether, these results show that the administration of urea restored TMAO compartmentalisation in a rat model of ASD, resulting in the normalisation of biochemical parameters and broad improvement in behaviour.

### Restoration of osmolarity/natraemia by urea in adult DHS rats

As ASD is not restricted to children, we also tested urea in three-month, that is adult DHS rats, which exhibited similar biochemical and behavioural alteration than young DHS rats (***Fig. 5a***). Urea normalised natremia (***Fig. 5b***) and markedly improved TMAO compartmentalisation (***Fig. 5c***). Plasma NAD^+^ levels (***Fig. 5d***) were restored to control values, but whole-blood 5-HT (***Fig. 5e***) and plasma melatonin levels (***Fig. 5f***) were only partially yet significantly restored toward normal values. From a behavioural standpoint, urea significantly improved most parameters (***Fig. 5g-m***). However, only sociability (***Fig. 5g-h***) and general locomotion (***Fig. 5k***) improved by more than 30%, while the improvement of other parameters remained physiologically marginal (***Fig. 5i-j*** and ***5l-m***). Altogether, these results show that the administration of urea mainly restored TMAO compartmentalisation in an adult rat model of ASD, although its impact on biochemical and behavioural parameters was less remarkable than in young animals.

**Fig. 5:**
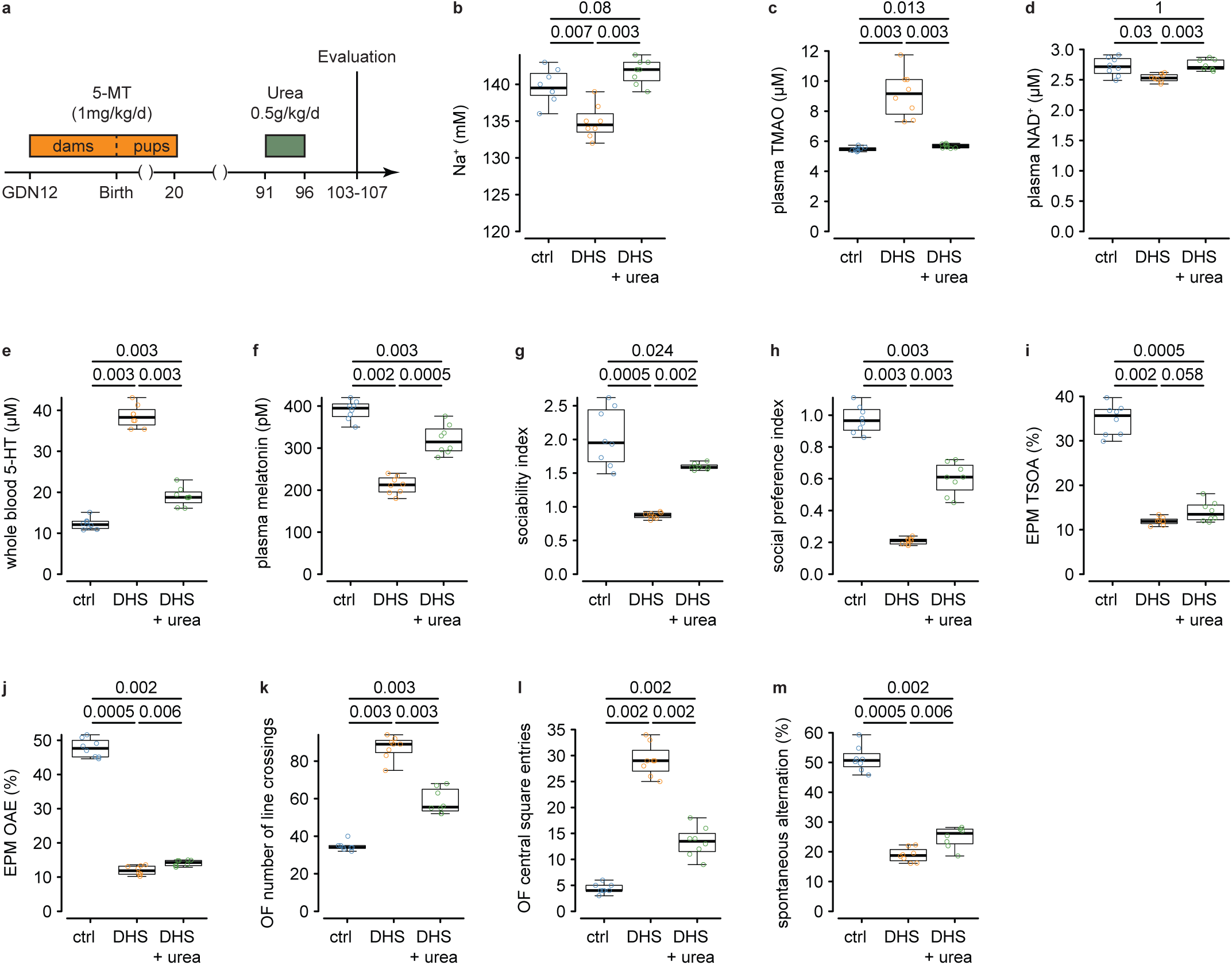
Effects of urea on adult DHS rats. **a,** schematic representation of the experimental protocol. Natremia (**b**), plasma TMAO (**c**), NAD^+^ (**d**) whole-blood serotonin (5-HT, **e**), plasma melatonin (**f**) concentrations in control rats (n = 8, blue), DHS rats (n = 8, orange) and DHS rats treated with urea at PDN91-PDN95 (n = 8, green). Sociability index (**g**), social preference index (**h**), time spent in the open arm (TSOA, **i**) and the number of open arm entries (OAE, **j**) in the elevated plus maze (EPM), the number of line crossing (**k**) and the number of central square entry (**l**) in the open field (OF), and the % of spontaneous alternation (**m**) in control rats (n = 8, blue), DHS rats (n = 8, orange) and DHS rats treated with urea (n = 8, green). Intergroup comparisons were performed using the Kruskal-Wallis test followed by the Wilcoxon sum-rank test corrected for multiple comparisons.

## Discussion

This study shows that TMAO miscompartmentalization appears to drive most biochemical and behavioural alterations in ASD. TMAO miscompartmentalization is likely driven by blood hypoosmolarity that is actionable pharmacologically, and we established the proof of concept that urea corrected TMAO miscompartmentalization and improved significantly biochemical and behavioural impairments in ASD. Overall, this study posits intracellular TMAO deficit as a common and major component of ASD pathophysiology and cellular physiology in general.

From a pathophysiological standpoint, lower intracellular TMAO levels observed in all individuals with ASD translated into an increase in plasma TMAO levels. Such elevation in TMAO plasma levels has been already described in ASD^25^, strongly suggesting that these findings are not restricted to our cohort. TMAO miscompartmentalization is correlated with hyponatraemia, although *ex vivo* data and previously published data^29,30^ suggest that hypoosmolarity is responsible for TMAO miscompartmentalization. Although all these measurements were performed in the blood, experimental hypotonic hyponatraemia caused cerebrospinal osmotic imbalance^39^, strongly suggesting that osmolarity is also altered in the brain of individuals with ASD and is in line with the higher extra-axial cerebrospinal fluid (CSF) volume in ASD^40^. Therefore, it is likely that peripheral TMAO miscompartmentalization is accompanied by central TMAO miscompartmentalization.

Our findings show that ASD is characterised by alterations in fluid/electrolyte homeostasis which provokes TMAO miscompartmentalization, hence physiological imbalance. Yet, TMAO miscompartmentalization is not *per se* causative of ASD, as patients with primary trimethylaminuria (*FMO3* mutation) do not exhibit any ASD-like phenotype. However, TMAO intracellular depletion becomes critical when, for instance, the serotonergic system is previously overstimulated by 5-MT (DHS rats). Interestingly, depletion or enhancement of 5-HT signalling is associated with ASD^41^, suggesting that any alteration of the serotonergic system could be a common cause of ASD, especially in idiopathic cases.

Therefore, we speculate that various mechanisms provoke the deregulation of the serotonergic system, which in turn causes alteration in brain development and maturation, along with an impairment in fluid/electrolyte homeostasis; the latter possibly resulting in alterations of the development/maturation of the hypothalamus/pituitary/adrenal axis (***Fig. 6***). Consequently, TMAO is miscompartmentalized, resulting in the alteration of numerous enzymatic activities that affect various metabolisms that contribute to most of the ASD-related and peripheral symptoms. As the correction of TMAO miscompartmentalization by urea improved all behavioural symptoms of ASD, we propose that the initial deregulation of the serotonergic system in itself may have mild behavioural consequences and that positive and negative reinforcements improve or worsen the initial symptoms, TMAO miscompartmentalization being a major negative reinforcement (**Fig. 6**).

**Fig. 6:**
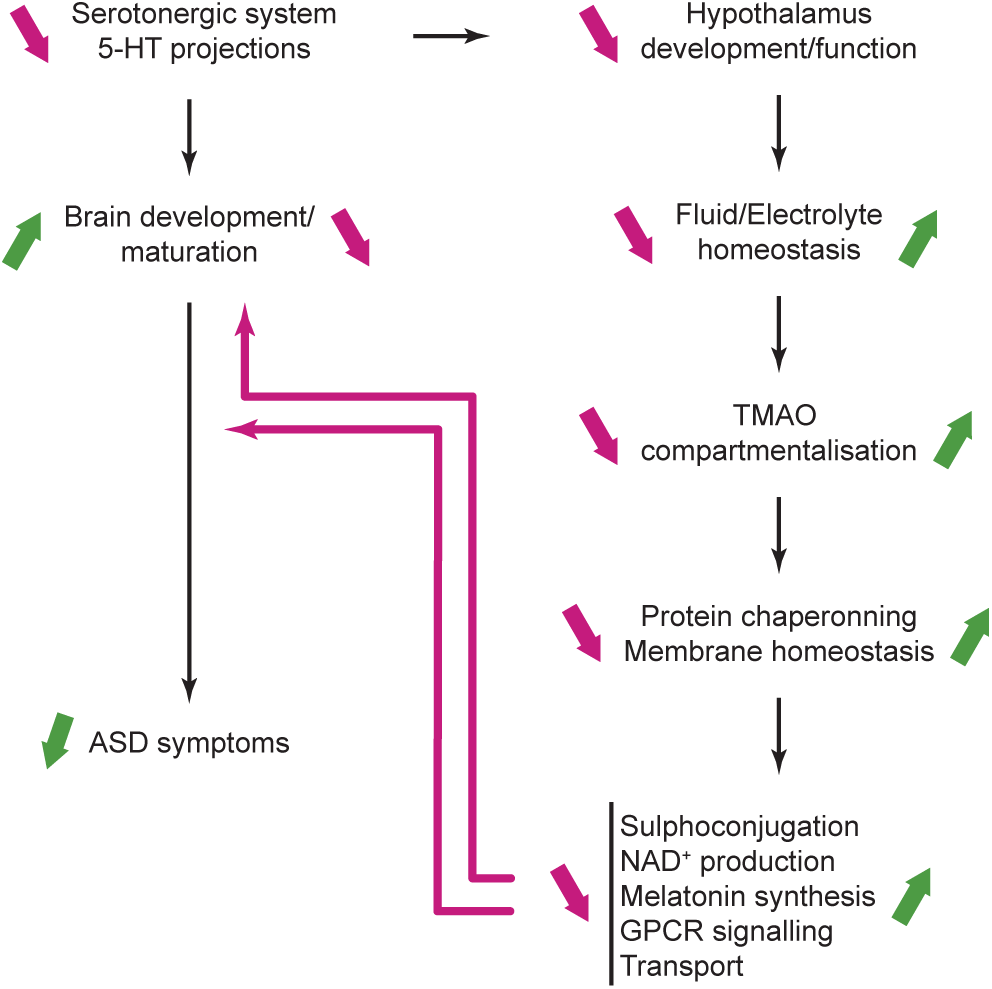
Tentative model involving osmolarity and inappropriate TMAO compartmentalisation in the pathophysiology of ASD. On the model, magenta indicates ASD and green the effect of urea.

From a therapeutic perspective, osmolarity can be readily corrected by dietary intake, such as salt^42^. During the sleep period, however, dietary intake is non-existent, resulting in water and salt losses only. This suggests that the negative impact of TMAO miscompartmentalization resets every sleep cycle, which prompts a therapeutic intervention to correct fluid/electrolyte imbalance appropriately. In that regard, urea is a salt-sparing diuretic that is clinically approved for the acute^37^ and chronic^38^ treatment of hyponatremia.

Consequently, urea provokes hemoconcentration capable of restoring osmolarity and TMAO compartmentalisation. Administration of urea showed major benefits in young DHS rats, whose age equivalence in humans (2-3 years old)^43^ conveniently fits within the time frame of ASD diagnosis^44^. Importantly, DHS rats shared strong behavioural and phenotypic resemblance with the individuals with ASD from our cohort, suggesting hope for transferability to humans. However, the phenotypic reversal observed after urea withdrawal advocates for chronic treatment and indicates that urea only palliates to fluid/electrolyte imbalance and not its cause. In adult rats, the results were less spectacular, possibly due to differences in neuroplasticity^43^ or hysteresis, which suggest that restoring urea treatment could improve post-diagnosis cerebral development in children with ASD. Nevertheless, the biochemical improvements observed in adult rats treated with urea still warrant trials in adults with ASD to alleviate the biochemical burden ASD is associated with. Interestingly, urea is not the first diuretic to be tested in ASD. The loop diuretic bumetanide has shown promising results^45,46^ for the treatment of ASD, although a recent phase III clinical trial failed to show efficacy^47^. Loop diuretics provoke the elimination of water and salts, with water loss exceeding that of salt, only resulting in moderate hemoconcentration. Nevertheless, these data reinforce hypoosmolarity as a therapeutic target in ASD and posit urea as a better diuretic candidate than loop diuretic to treat ASD. Finally, these results suggest that loop diuretics act via a partial restoration of fluid/electrolyte homeostasis rather than a direct improvement in GABAergic signalling.

More broadly, our results show that TMAO is important, yet not essential, in cell physiology. Based on our data, TMAO appears to chaperone homodimeric interactions (sulphotransferase activities) and protein complexes (melatonin synthesis) and play a major role in membrane fluidity (AVP binding), hence GPCR signalling (V1aR) and transport (OCT-2), under physiological conditions. However, it should be noted that TMAO does not always have a positive effect on proteins^48,49^, indicating that TMAO homeostasis is important. A recent study has shown that only ∼5% of cellular water content was free and suggested that a main impact of change in electric charge by post-translational modification was to modify the behaviour of water molecules around a protein^50^. Importantly, TMAO does not physically interact with either proteins^51–53^ or lipids^20,21,54^ but rather interacts with the surrounding water molecules^53^. Furthermore, the high TMAO dipole moment modifies water distribution around it^55^. Therefore, given the ubiquitous nature of TMAO-water interactions, we anticipate TMAO to impact a vast majority of proteins and lipids in the cell.

In conclusion, TMAO appears as a major modulator of cell physiology, which, when combined with neurodevelopmental alteration, acts as a major worsening factor in ASD. Given the phenotypic similarities between DHS rats and individuals with ASD, urea offers a promising therapeutic option for ASD that warrants clinical trials in humans.

## Acknowledgements

The authors would like to thank Professors Marion Leboyer and Richard Delorme for providing human biological samples, and Dr Michel Vodovar for critical discussions. This work is dedicated to the memory of Mosé Da Prada (1930-1995), whose pioneering interest in osmolarity laid the foundation for this work. We thank The Warning and Foo Fighters for inspiration while writing this manuscript. This work was supported by funding from Institut National de la Santé et Recherche Médicale and Université Paris Cité.

## Author contributions

JML and NV equally contributed to this study.

## Competing interests

The authors have no competing interests to declare.

## Notes

### Competing Interest Statement

The authors have declared no competing interest.

### Summary of Updates

In Figure 3,4, and 5, mthe P-values presented corresponded to the raw P-values not corrected for multiple comparisons (holm). The new figure now display the corrected P-value. This correction did not impact the statistical significance of the results nor did it change the interpretation of the results.

## References

1. American Psychiatric Association. Diagnostic and statistical manual of mental disorders (5th ed.). (2013).

2 Miles, J. H. Autism spectrum disorders--a genetics review. Genet Med 13, 278–294, doi:10.1097/GIM.0b013e3181ff67ba (2011).

3 Doi, M., Li, M., Usui, N. & Shimada, S. Genomic Strategies for Understanding the Pathophysiology of Autism Spectrum Disorder. Front Mol Neurosci 15, 930941, doi:10.3389/fnmol.2022.930941 (2022).

4 Rolland, T. et al. Phenotypic effects of genetic variants associated with autism. Nat Med 29, 1671–1680, doi:10.1038/s41591-023-02408-2 (2023).

5 Safadi, J. M., Quinton, A. M. G., Lennox, B. R., Burnet, P. W. J. & Minichino, A. Gut dysbiosis in severe mental illness and chronic fatigue: a novel trans-diagnostic construct? A systematic review and meta-analysis. Mol Psychiatry 27, 141–153, doi:10.1038/s41380-021-01032-1 (2022).

6 Yap, C. X. et al. Autism-related dietary preferences mediate autism-gut microbiome associations. Cell 184, 5916–5931 e5917, doi:10.1016/j.cell.2021.10.015 (2021).

7 Pagan, C. et al. Decreased phenol sulfotransferase activities associated with hyperserotonemia in autism spectrum disorders. Transl Psychiatry 11, 23, doi:10.1038/s41398-020-01125-5 (2021).

8 Pagan, C. et al. Disruption of melatonin synthesis is associated with impaired 14-3-3 and miR-451 levels in patients with autism spectrum disorders. Sci Rep 7, 2096, doi:10.1038/s41598-017-02152-x (2017).

9 Launay, J. M. et al. Impact of IDO activation and alterations in the kynurenine pathway on hyperserotonemia, NAD(+) production, and AhR activation in autism spectrum disorder. Transl Psychiatry 13, 380, doi:10.1038/s41398-023-02687-w (2023).

10 Loo, R. L., Chan, Q., Nicholson, J. K. & Holmes, E. Balancing the Equation: A Natural History of Trimethylamine and Trimethylamine-N-oxide. J Proteome Res 21, 560–589, doi:10.1021/acs.jproteome.1c00851 (2022).

11 Fahnert, B. Using folding promoting agents in recombinant protein production: a review. Methods Mol Biol 824, 3–36, doi:10.1007/978-1-61779-433-9_1 (2012).

12 Cho, I. C. & Swaisgood, H. Factors affecting tetramer dissociation of rabbit muscle lactate dehydrogenase and reactivity of its sulfhydryl groups. Biochemistry 12, 1572–1577, doi:10.1021/bi00732a017 (1973).

13 Kumari, K., Singh, K. S., Singh, K., Bakhshi, R. & Singh, L. R. TMAO to the rescue of pathogenic protein variants. Biochim Biophys Acta Gen Subj 1866, 130214, doi:10.1016/j.bbagen.2022.130214 (2022).

14 Miller, A. L. et al. Restored mutant receptor:Corticoid binding in chaperone complexes by trimethylamine N-oxide. PLoS One 12, e0174183, doi:10.1371/journal.pone.0174183 (2017).

15 Yancey, P. H. & Somero, G. N. Counteraction of urea destabilization of protein structure by methylamine osmoregulatory compounds of elasmobranch fishes. Biochem J 183, 317–323, doi:10.1042/bj1830317 (1979).

16 Yancey, P. H. & Siebenaller, J. F. Trimethylamine oxide stabilizes teleost and mammalian lactate dehydrogenases against inactivation by hydrostatic pressure and trypsinolysis. J Exp Biol 202, 3597–3603, doi:10.1242/jeb.202.24.3597 (1999).

17 Lambert, D., Leipply, D. & Draper, D. E. The osmolyte TMAO stabilizes native RNA tertiary structures in the absence of Mg2+: evidence for a large barrier to folding from phosphate dehydration. J Mol Biol 404, 138–157, doi:10.1016/j.jmb.2010.09.043 (2010).

18 Nandi, S., Pyne, A., Layek, S., Arora, C. & Sarkar, N. The Dietary Nutrient Trimethylamine N-Oxide Affects the Phospholipid Vesicle Membrane: Probable Route to Adverse Intake. J Phys Chem Lett 12, 12411–12418, doi:10.1021/acs.jpclett.1c03201 (2021).

19 Higgins, T., Chaykin, S., Hammond, K. B. & Humbert, J. R. Trimethylamine N-oxide synthesis: a human variant. Biochem Med 6, 392–396, doi:10.1016/0006-2944(72)90025-7 (1972).

20 Maiti, A. & Daschakraborty, S. Effect of TMAO on the Structure and Phase Transition of Lipid Membranes: Potential Role of TMAO in Stabilizing Cell Membranes under Osmotic Stress. J Phys Chem B 125, 1167–1180, doi:10.1021/acs.jpcb.0c08335 (2021).

21 Manisegaran, M., Bornemann, S., Kiesel, I. & Winter, R. Effects of the deep-sea osmolyte TMAO on the temperature and pressure dependent structure and phase behavior of lipid membranes. Phys Chem Chem Phys 21, 18533–18540, doi:10.1039/c9cp03812d (2019).

22 Koeth, R. A. et al. Intestinal microbiota metabolism of L-carnitine, a nutrient in red meat, promotes atherosclerosis. Nat Med 19, 576–585, doi:10.1038/nm.3145 (2013).

23 Tang, W. H. et al. Intestinal microbial metabolism of phosphatidylcholine and cardiovascular risk. N Engl J Med 368, 1575–1584, doi:10.1056/NEJMoa1109400 (2013).

24 Zhu, W. et al. Gut Microbial Metabolite TMAO Enhances Platelet Hyperreactivity and Thrombosis Risk. Cell 165, 111–124, doi:10.1016/j.cell.2016.02.011 (2016).

25 Quan, L. et al. Plasma trimethylamine N-oxide, a gut microbe-generated phosphatidylcholine metabolite, is associated with autism spectrum disorders. Neurotoxicology 76, 93–98, doi:10.1016/j.neuro.2019.10.012 (2020).

26 Obsil, T., Ghirlando, R., Klein, D. C., Ganguly, S. & Dyda, F. Crystal structure of the 14-3-3zeta:serotonin N-acetyltransferase complex. a role for scaffolding in enzyme regulation. Cell 105, 257–267, doi:10.1016/s0092-8674(01)00316-6 (2001).

27 Li, H., Nguyen, H. H., Ogorzalek Loo, R. R., Campuzano, I. D. G. & Loo, J. A. An integrated native mass spectrometry and top-down proteomics method that connects sequence to structure and function of macromolecular complexes. Nat Chem 10, 139–148, doi:10.1038/nchem.2908 (2018).

28 Pagan, C. et al. The serotonin-N-acetylserotonin-melatonin pathway as a biomarker for autism spectrum disorders. Transl Psychiatry 4, e479, doi:10.1038/tp.2014.120 (2014).

29 Koomoa, D. L., Musch, M. W., MacLean, A. V. & Goldstein, L. Volume-activated trimethylamine oxide efflux in red blood cells of spiny dogfish (Squalus acanthias). Am J Physiol Regul Integr Comp Physiol 281, R803–810, doi:10.1152/ajpregu.2001.281.3.R803 (2001).

30 MacLellan, R. J. et al. Chaperone roles for TMAO and HSP70 during hyposmotic stress in the spiny dogfish shark (Squalus acanthias). J Comp Physiol B 185, 729–740, doi:10.1007/s00360-015-0916-6 (2015).

31 Teft, W. A. et al. Identification and Characterization of Trimethylamine-N-oxide Uptake and Efflux Transporters. Mol Pharm 14, 310–318, doi:10.1021/acs.molpharmaceut.6b00937 (2017).

32 McNamara, I. M., Borella, A. W., Bialowas, L. A. & Whitaker-Azmitia, P. M. Further studies in the developmental hyperserotonemia model (DHS) of autism: social, behavioral and peptide changes. Brain Res 1189, 203–214, doi:10.1016/j.brainres.2007.10.063 (2008).

33 Hormann, S., Gai, Z., Kullak-Ublick, G. A. & Visentin, M. Plasma Membrane Cholesterol Regulates the Allosteric Binding of 1-Methyl-4-Phenylpyridinium to Organic Cation Transporter 2 (SLC22A2). J Pharmacol Exp Ther 372, 46–53, doi:10.1124/jpet.119.260877 (2020).

34 Wiegand, V. & Gimpl, G. Specification of the cholesterol interaction with the oxytocin receptor using a chimeric receptor approach. Eur J Pharmacol 676, 12–19, doi:10.1016/j.ejphar.2011.11.041 (2012).

35 Parker, K. J. et al. A randomized placebo-controlled pilot trial shows that intranasal vasopressin improves social deficits in children with autism. Sci Transl Med 11, doi:10.1126/scitranslmed.aau7356 (2019).

36 Launay, J. M. et al. V1a-vasopressin specific receptors on human platelets: potentiation by ADP and epinephrine and evidence for homologous down-regulation. Thromb Res 45, 323–331, doi:10.1016/0049-3848(87)90221-0 (1987).

37 Decaux, G. et al. Actual Therapeutic Indication of an Old Drug: Urea for Treatment of Severely Symptomatic and Mild Chronic Hyponatremia Related to SIADH. J Clin Med 3, 1043–1049, doi:10.3390/jcm3031043 (2014).

38 Rondon-Berrios, H. Urea for Chronic Hyponatremia. Blood Purif 49, 212–218, doi:10.1159/000503773 (2020).

39 Melton, J. E. & Nattie, E. E. Brain and CSF water and ions during dilutional and isosmotic hyponatremia in the rat. Am J Physiol 244, R724–732, doi:10.1152/ajpregu.1983.244.5.R724 (1983).

40 Shen, M. D. et al. Extra-axial cerebrospinal fluid in high-risk and normal-risk children with autism aged 2-4 years: a case-control study. Lancet Psychiatry 5, 895–904, doi:10.1016/S2215-0366(18)30294-3 (2018).

41 Garbarino, V. R., Gilman, T. L., Daws, L. C. & Gould, G. G. Extreme enhancement or depletion of serotonin transporter function and serotonin availability in autism spectrum disorder. Pharmacol Res 140, 85–99, doi:10.1016/j.phrs.2018.07.010 (2019).

42 Good, P. Do salt cravings in children with autistic disorders reveal low blood sodium depleting brain taurine and glutamine? Med Hypotheses 77, 1015–1021, doi:10.1016/j.mehy.2011.08.038 (2011).

43 Semple, B. D., Blomgren, K., Gimlin, K., Ferriero, D. M. & Noble-Haeusslein, L. J. Brain development in rodents and humans: Identifying benchmarks of maturation and vulnerability to injury across species. Prog Neurobiol 106-107, 1–16, doi:10.1016/j.pneurobio.2013.04.001 (2013).

44 Hyman, S. L., Levy, S. E., Myers, S. M., Council On Children With Disabilities, S. O. D. & Behavioral, P. Identification, Evaluation, and Management of Children With Autism Spectrum Disorder. Pediatrics 145, doi:10.1542/peds.2019-3447 (2020).

45 Lemonnier, E. et al. Effects of bumetanide on neurobehavioral function in children and adolescents with autism spectrum disorders. Transl Psychiatry 7, e1056, doi:10.1038/tp.2017.10 (2017).

46 Sprengers, J. J. et al. Bumetanide for Core Symptoms of Autism Spectrum Disorder (BAMBI): A Single Center, Double-Blinded, Participant-Randomized, Placebo-Controlled, Phase-2 Superiority Trial. J Am Acad Child Adolesc Psychiatry 60, 865–876, doi:10.1016/j.jaac.2020.07.888 (2021).

47 Fuentes, J. et al. Bumetanide oral solution for the treatment of children and adolescents with autism spectrum disorder: Results from two randomized phase III studies. Autism Res 16, 2021–2034, doi:10.1002/aur.3005 (2023).

48 Buawangpong, N., Pinyopornpanish, K., Siri-Angkul, N., Chattipakorn, N. & Chattipakorn, S. C. The role of trimethylamine-N-Oxide in the development of Alzheimer’s disease. J Cell Physiol 237, 1661–1685, doi:10.1002/jcp.30646 (2022).

49 Sackett, D. L. Natural osmolyte trimethylamine N-oxide stimulates tubulin polymerization and reverses urea inhibition. Am J Physiol 273, R669–676, doi:10.1152/ajpregu.1997.273.2.R669 (1997).

50 Watson, J. L. et al. Macromolecular condensation buffers intracellular water potential. Nature 623, 842–852, doi:10.1038/s41586-023-06626-z (2023).

51 Zou, Q., Bennion, B. J., Daggett, V. & Murphy, K. P. The molecular mechanism of stabilization of proteins by TMAO and its ability to counteract the effects of urea. J Am Chem Soc 124, 1192–1202, doi:10.1021/ja004206b (2002).

52 Hu, C. Y., Lynch, G. C., Kokubo, H. & Pettitt, B. M. Trimethylamine N-oxide influence on the backbone of proteins: an oligoglycine model. Proteins 78, 695–704, doi:10.1002/prot.22598 (2010).

53 Canchi, D. R., Jayasimha, P., Rau, D. C., Makhatadze, G. I. & Garcia, A. E. Molecular mechanism for the preferential exclusion of TMAO from protein surfaces. J Phys Chem B 116, 12095–12104, doi:10.1021/jp304298c (2012).

54 Sukenik, S., Dunsky, S., Barnoy, A., Shumilin, I. & Harries, D. TMAO mediates effective attraction between lipid membranes by partitioning unevenly between bulk and lipid domains. Phys Chem Chem Phys 19, 29862–29871, doi:10.1039/c7cp04603k (2017).

55 Kuroki, N., Uchino, Y., Funakura, T. & Mori, H. Electronic fluctuation difference between trimethylamine N-oxide and tert-butyl alcohol in water. Sci Rep 12, 19417, doi:10.1038/s41598-022-24049-0 (2022).

