## Supplementary material for "TMAO miscompartmentalization is a reversible driver of autism pathophysiology": Methods and Supplementary Figures

**Supplementary Data**

Materials and methods

Supplementary Figures 1-3

### Methods

#### *Ethics statement*

This study was approved by the local ethics committee (Comité de Protection des Personnes Ile de France IX) and was performed according to the current revision of the Helsinki Declaration. Animal care and experimental procedures complied with the European Communities Council Directive (CEE 86/609/EEC), EU Directive 2017/32/EU, and French Departmental Direction of Animal Protection (2019-11 #1027).

#### *Human cohort*

The cohort used has been already described[7,9]. It consisted of 90 individuals with ASD and 106 neurotypic controls. Written informed consent was obtained after oral and written information from all participants, their parents or legal guardians. Clinical evaluations and blood sampling of individuals with ASD (diagnosed according to DSM-IVTR) and control individuals from the general population investigated for genetics and blood biochemistry have previously been detailed previously[7,28]. Venous blood samples were collected after a 48-hour diet that excluded important sources of serotonin and tryptophan. Fasting blood samples were collected into tubes containing 109 mM sodium citrate (Greiner Bio-One) with a 9:1 blood-to-anticoagulant ratio. After removing 0.5mL of whole blood (for serotonin measurement), the remainder was centrifuged at 1,000 x g for 10 minutes, and the supernatant was then centrifuged at 3,000 x g for 15 minutes to isolate platelets and obtain platelet-poor-plasma. Importantly, the time between blood sampling and platelet/plasma isolation was less than 1 hour. Whole blood, platelets, and plasmas were aliquoted into barcode-labelled cryovials and stored at -80°C until investigations.

*Natraemia*

Natraemia was measured by potentiometry on an Abbott Architect c8000 (Abbott Laboratories, Abbott Park, IL, USA).

*Osmolarity experiments*

For osmolarity experiments, human platelets from healthy donors were isolated after blood collection on pre-chilled vacutainer tubes containing sodium citrate. Blood samples were centrifuged at 1000×g/10 min, and the supernatant was further centrifuged at 3000×g/10 min. The platelet pellet was resuspended in a Tyrode-Tris buffer, pH 7.40, 290-310 mOsm/L (NaCl 130 mM; KCl 5.6 mM; Tris 12.4 mM; $Na_4EDTA$ 2.1 mM; $NaH_2PO_4$ 0.9 mM; sucrose 13.1 mM; and dextrose 11.1 mM) which was selected as the most efficient for the study of 5-HT uptake and platelet physiology on isolated platelets[56]. Tyrode solution was diluted with water to decrease osmolarity to 285 and 275 mOsm/L.

Immunoprecipitation/Mass spectrometry

Platelet samples from healthy individuals were obtained as described above. Platelet lysis was achieved by -SH-activated toxin treatment[57]. Alveolysin was purified to homogeneity and 150 haemolytic units per $1.5x10^9$ platelets (equivalent to about 375 ng or 6 pmol of protein) were added (10:1) to platelet suspensions at 4°C and the suspensions were warmed to 37°C for enzyme measurement. This amount of toxin binds to platelets at 4°C and elicits complete lysis at 37°C since about 16 molecules of toxin are sufficient to lyse one human platelet[58]. Immunoprecipitations were performed using human anti-p31T-AANAT (ref: AB-5467, Merck, Darmstadt, Germany) or anti-ASMT (ref: A72945, EpiGentek, Farmingdale, NY, USA). GC-MS analysis was carried out on a TRACE GC 2000 Series (ThermoQuest CE Instruments, Austin TX, USA) gas chromatograph, interfaced with GCQ Plus (ThermoQuest) mass detector with ion trap analyser, operating in EI mode (70 eV). The capillary GC column was an OV-1 (30 m x 0.25 mm ID, OV-1 bonded, 0.25 µm film thickness ref 1105-2502 105, Ohio Valley Specialty Co, Marietta OH, USA). Helium was the carrier gas at a flow rate of 1.0 mL/min. A temperature program was adopted: initial temperature was 40°C (hold time: 5 min), then ramped by 10°C/min to 220°C (hold time: 5 min). The temperatures of transfer line and ionization source were maintained at 250 and 200°C respectively. The GC was operated in splitless mode; the injector base temperature was set at 250°C. The mass spectra were recorded in full scan mode (30–200 amu) to collect the total ion current (TIC) chromatograms. Quantitation was carried out by using the extracted ion chromatograms (XIC) by selecting qualifier and quantifier fragment ions of the studied analytes. Standard TMAO and the concentrated eluates were run under the same conditions.

*TMAO measurements*

Twenty-five µL of plasma or platelet lysate and 300 µL of acetonitrile:methanol:water (5:4:1; v:v:v) containing the internal standard (IS) d9-TMAO for quantification were mixed and vigorously vortexed for 20 s. The samples were re-equilibrated on ice for 30 min and centrifuged for 10 min at 25,100×g at 4°C. The supernatant was then transferred into a specific vial before liquid chromatography-mass spectrometry (LC-MS) analysis. Matrix-matched calibration curves were generated using a human control plasma pool spiked with the standards. The concentration range of the calibration curves was 0–250 µM. An ultra-high-performance LC system coupled with a 6490 triple-quadrupole mass spectrometer (QqQ, Agilent Technologies) using an electrospray as an ion source (LC-ESI-QqQ) working in positive mode was used to analyse the extracts. An ACQUITY UPLC BEHHILIC column (1.7 mm, 2.1 × 150 mm, Waters) and a gradient mobile phase containing water with 50 mM ammonium acetate (phase A) and acetonitrile (phase B) were used to perform the chromatographic separation. The gradient was as follows: for 0.5 min, isocratic at 75% B; from 0.5 to 2 min, reduced to 65% B; from 2 to 2.1 min, decreased to 45% B; from 2.1 to 3.9

min, maintained at 45% B; from 3.9 to 4 min, hiked up to 75% B; and, finally, until 5.5 min, the column was equilibrated at 75% B. The flow throughout this process was 0.6 mL/min. Two hundred microliters of plasma extract were injected into the LC system. The mass spectrometer conditions were: a drying gas temperature of 280°C and a sheath gas temperature of 400°C; source and sheath gas flows of 20 and 12 L/min, respectively; a nebulizer flow of 60 psi; a nozzle voltage of 500 V; a capillary voltage of 2500 V; and iFunnel and HRF values of 110 and 80 V, respectively. The QqQ worked in the multiple reaction monitoring (MRM) mode and used predetermined transitions and collision energies (CE(V)) of 16 (transition 76 → 58) and 8 (transition 76 → 59) for TMAO, and of 16 (transition 85 → 67) and 8 (transition 85 → 68) for d9-TMAO (IS) respectively.

*Other biochemical measurements*

Serotonin and kynurenine metabolites were measured by LC-MS/MS as previously described[59]. Melatonin was measured using a radioimmunoassay (ref: RK-MEL2, Novolytix, Switzerland) according to the manufacturer's instructions. $NAD^+$ concentrations were determined by LC-MS as described previously[60]. The amount of 14-3-3 proteins was determined using the commercial 14-3-3 Pro ELISA kit from MyBioSource (San Diego, CA, USA). The immunogen used to produce the antibodies is recombinant full-length human 14-3-3 γ. The kit has about 40% cross-reactivity with 14-3-3 ζ, ε, σ, τ (MyBioSource, personal communication).

*Enzymatic activities*

Platelet or tissue lysis was achieved by -SH-activated toxin treatment[57] as described above. Separate experiments showed that the addition of equivalent amounts of toxin to cells already disrupted by homogenization produced no change in the rates of enzyme activities measured at least in duplicate. Under the conditions employed here, enzyme rates were linear with time and with platelet protein concentrations ranging from 0.01 to 0.1 mg of protein per mL.

PST (EC 2.8.2.1) activities were determined by radioenzymology using p-nitrophenol as substrates for humans (PST-P) and mice (PST)[7].

Measurement of AANAT activity[61] was performed at 37°C in 1.5 mL polypropylene microtubes. The reaction was started by the addition of enzyme preparation (25 µL) to 40 µL buffer containing 20 mM sodium citrate (pH 7.00), 300 mM NaCl, 1 mM EDTA, 0.05 mg/ml BSA, 50 nM [acetyl-$^3$H]-CoA, and 500 nM tryptamine. After 20 min the reaction was stopped by adding 1 mL of chloroform and vortexing the mixture. The reaction product, N-[$^3$H]-acetyl tryptamine, was extracted into the chloroform which was then washed once with 0.1 mL of 0.1 mM sodium phosphate buffer, pH 6.80. A 500 µL aliquot of the chloroform was then transferred to a scintillation vial. Three mL of scintillation fluid were added and the radioactivity was measured by a liquid scintillation counter (Beckman LS 6000SC).

The reaction mixtures for ASMT assay[62] contained in a final volume of 50 µL the following: N-acetylserotonin (1 mM), tissue homogenate (25 µL), and 0.1 mM S-[methyl-$^3$H]-adenosyl-L-methionine. Reactions were carried out in plastic Eppendorf tubes at 37°C for 30 min in 50 mM sodium phosphate buffer (pH 7.90). Reactions were stopped by the addition of 100 µL of 0.5 M borate buffer (pH 10.50) to the tubes. Blanks consisted of reaction mixtures minus N-acetylserotonin. One mL of chloroform was added. The mixture was vortexed twice for 15 s to extract the product, [$^3$H]-melatonin, into the chloroform. The organic layer was washed once with borate buffer, and an aliquot (500 µL) was dried and its radioactivity measured. To verify that the activities measured in platelets reflect authentic AANAT and ASMT and not non-specific actions of other N-acetyltransferases and methyltransferases, si-RNA specific for each enzyme (SR300008 and SR300317, Origene; 1 nM final concentrations after transfection by the siTran 1.0 as recommended by Origene) were used. 3-HAO activity was determined by the decrease in 3-HK measured by LC-MS/MS[59] in 10 mM Hepes, pH 6.5, containing 0.3 mM $Fe(NH_4SO_4)_2$.

*In vitro enzymatic reactions*

Human recombinant AANAT, ASMT and 14-3-3ζ were purified as previously described[63-65] and mixed in 1:1:2 ratio in the presence of 10 nM 5-HT binoxalate in optimal conditions for melatonin production[66]. Human recombinant *SULT1A1* gene product (PST-P) was expressed in budding yeast[67] and purified as previously described[68]. PST-P activity was measured using p-nitrophenol as substrate[7]. Human recombinant 3-HAO was purified as previously described[69] and its activity measured as described above.

*Binding and uptake experiments*

Platelet AVP and D-22 bindings were performed as previously published[70,71] with 5 nM $[^3H]$AVP and 7.5 nM $[^3H]$D-22 respectively. Platelet $IP_3$ concentrations of were measured radio-immunologically as described previously[72] except the TRK 1000 kit (Amersham) was replaced by the NEK 064 kit (Perkin-Elmer, Boston, MA, USA). The recovery was 90.1 ± 4.8 % and the intra- and inter-assay coefficients of variations were 4.4% and 7%, respectively. Platelet $MPP^+$ uptake ($[^3H]MPP^+$ ref RCTT0970 from Tritec, Teufen, Switzerland) were performed as previously published[73].

*Mice*

Wild-type male FVB/N were from Charles River and housed in light- (on from 08:00 – 20:00 h) and temperature (22°C)-controlled testing rooms, with food and water available *ad libitum*. Blood samples were obtained from the submandibular (facial) vein at baseline, after administration of 50µg/kg TMAO, and after administration of 3mg/kg of the FMO3 inhibitor phenylthiourea. The same animals were used throughout the experiment. A one-week wash-out period was allowed between the administration of exogenous TMAO and FMO3i phenylthiourea, once confirmed that plasma TMAO levels returned to baseline levels.

*Rats*

The developmental hyperserotonaemia (DHS) rat model for ASD was performed as previously described[32]. Nulliparous Sprague–Dawley female rats (180-220 g) were from Charles River and housed in light- (on from 08:00 – 20:00 h) and temperature (22°C)-controlled testing rooms, with food and water available *ad libitum*. They were mated overnight, and fertilization was determined using a vaginal smear; this day was considered gestational day 0 (GD 0). Eight pregnant females were administered subcutaneously a single dose (1.0 mg/ kg of body weight) of 5-methoxytryptamine (5-MT, Sigma Aldrich St. Louis MO) daily from gestational day (GD )12 to parturition and five pregnant females received the vehicle in equal volumes on the same days. Upon parturition (PND 0), litters were culled to five males and five females. Male pups exposed to 5-MT *in utero* were further administered 5-MT from PND0 to PND20, to induce DHS. Similarly, pups only exposed to the vehicle prenatally were administered the vehicle in equal volumes. At PND21 (young rats) or PND91 (adult rats), 0.5 g/kg body weight/day urea (Sigma-Aldrich) was administered subcutaneously using an osmotic pump (AZLET) for 5 days.

*Behavioural analyses*

All behavioural studies were conducted during the light phase, between 09:00 and 18:00 hours, and were assessed by three independent observers blinded for the intervention; each data point correspond to the median of the three independent measures (relative variation < 10%) . To assess the impact of urea on DHS rats, animals were exposed to behavioural protocols on open field-spontaneous locomotory activity (PND33, PND60, PND90 in young rats and PDN103 for adult rats)[74], three-chambered social interaction–social behaviour/interaction (PND34-35, PDN61-62, PDN91-92 for young rats and PDN104-105 for adult rats), y-maze–repetitive behaviour (PND36, PND63, PND93 and PDN106 for adult rats)[75] and elevated plus maze (EPM)–anxiety (PND37, PND64, PND94 for young rats and PND107 for adult rats)[76]. Animals were placed in the testing area 5 days before the beginning of the behavioural experiments. To reduce the chances of olfaction-related cues during testing, the test arena was cleaned with 70% v/v ethyl alcohol and dried between each consecutive trial. In young rats administered with urea from PDN21 to PDN25

*Locomotor activity:* Spontaneous increase in locomotion is extensively reported as one of the important features of the DHS model, this hyper-locomotion was measured using an open field apparatus. The apparatus measured 90 cm x 90 cm with 50 cm high walls made of dark-coloured wood. The animals (PND33, PND60, PND90 in young rats and PDN103 for adult rats) were introduced individually into the centre of the arena for a single 10-minute trial period[77]. The total number of line crossings and the total number of central square entries were recorded to assess changes in the locomotion of the animals.

*Social behaviour*

Diminished and abnormal social interaction is a core phenotype of ASD, which may be mimicked by the DHS model[78]. This change in social behaviour was assessed using the three-chambered social interaction test protocol (PND34-35, PDN61-62, PDN91-92 for young rats and PDN104-105 for adult rats)[75]. The test arena measured 76 cm x 30 cm x 35 cm and was divided into three chambers with an access point between each chamber. Animals had free access to all the chambers and each trial began with the animal being placed in the central chamber. To encourage exploration of the side chambers, all animals were habituated to the apparatus for 5 min before initiation of the test trial. After ending of the habituation period, the rats were tested in the sociability phase lasting for 10 min. Animals to be placed under a wire cage were habituated to the wire cage for 30 min, 24 hrs before the beginning of the sociability phase. In the sociability phase, a stranger animal was placed under the wired cage in either (left or right) side chamber, while in the other chamber, an empty cage would be placed. To avoid side preferences, the placement of wired cages was randomized and the chamber with the stranger animal and the empty cage are referred to as the stranger chamber and the empty chamber, respectively. Upon conclusion of the sociability phase, the social preference phase was initiated 2 hrs after the last animal trial. In the social preference phase, each animal was allowed to explore the complete arena for 10 min. During this phase, the animal previously considered a stranger was now familiar and a new stranger animal was introduced into the previously empty cage. So, the two chambers now became the familiar chamber and the novel chamber. The time spent by test animals in both chambers containing the cages was measured. The sociability index (SI) and the social preference index (SPI) were calculated according to the following formula[76]: Sociability index = total time in the stranger chamber/total time in the empty chamber, and Sociability Preference Index (SPI) = total time in the novel chamber/total time in the familiar chamber, respectively.

*Anxiety:* Anxiety is the most common co-morbid trait expressed with ASD[79]. An elevated plus maze (EPM) is a commonly used apparatus to assess anxiety-like behaviour in animal models of ASD. The EPM apparatus was made up of wood with four arms at 90° to each other. Two open arms and two closed arms of 50 x 10 cm dimensions, enclosed by a 40 cm high wall. To facilitate exploration of the maze, all animals were placed in a pre-test arena for 5 min each (PND37, PND64, PND94 for young rats and PND107 for adult rats). Soon after, the animals were transferred to the EPM placed 50 cm high from the ground. All animals were released in the centre of the maze, pointing towards the open arm and entries along with the time spent in each arm were recorded for 5 min. The basis of this test is the conflict associated between the two parts of the maze *i.e.* the open arms which are aversive, bright and unprotected and the closed arms which are covered, shadowy and protected. To this, the number of open arm entries and the closed arm entries, time spent in the open arm and time spent in the closed arm were measured to finally calculate % open arm entries and % open arm time, using the following formula[80]: % open arm entries (OAE) = 100 x number of entries in open arm/(closed arm entries + open arm entries) and % time spent in open arm (TSOA) = 100 x time spent in open arm/(time spent in open arm + time spent in the closed arm)

*Repetitive behaviour:* One of the core diagnostic features of ASD is the presence of stereotypical or repetitive behaviour, this key clinical feature is replicated by inducing DHS[81]. The extent of % spontaneous alteration is regarded as a measure of stereotypy or repetitive behaviour in animals. A y-maze apparatus was used to assess % spontaneous alterations[82]. The maze makes a Y shape, with three arms of equal lengths each at an angle of 120° from the other. One of the arms was considered the start arm and all animals were placed at the end of this arm pointing towards the centre of the maze. The animals were subjected to 8 min of testing in the y-maze (PND36, PND63, PND93 and PDN106 for adult rats). The exploration of three different arms in succession was considered as one alternation. Serial arm entries were observed for each animal to calculate % spontaneous alternations. The % spontaneous alternation can be calculated with the following formula: % Spontaneous alternation = 100 x total alternations/(total arm entries−2)

*Statistical analysis*

All the statistical analyses were performed using R-statistical software (version 4.2.1). Data normality was assessed using the Shapiro-Wilk test. Intergroup comparisons were performed using the sum-rank Wilcoxon test or the Kruskal-Wallis test followed by the sum-rank Wilcoxon test corrected for multiple comparisons (holm). Longitudinal data were analysed using repeated-measure ANOVA on log-transformed data followed by the Tukey Honest Statistical Significance Difference test. Correlations between variables were evaluated using Spearman's comparison coefficient (ρ) or linear regression. A P-value < 0.05 was considered significant and exact P-values are reported.

### References


7 Pagan, C. *et al.* Decreased phenol sulfotransferase activities associated with hyperserotonemia in autism spectrum disorders. *Transl Psychiatry* **11**, 23, doi:10.1038/s41398-020-01125-5 (2021).

9 Launay, J. M. *et al.* Impact of IDO activation and alterations in the kynurenine pathway on hyperserotonemia, NAD(+) production, and AhR activation in autism spectrum disorder. *Transl Psychiatry* **13**, 380, doi:10.1038/s41398-023-02687-w (2023).

28 Pagan, C. *et al.* The serotonin-N-acetylserotonin-melatonin pathway as a biomarker for autism spectrum disorders. *Transl Psychiatry* **4**, e479, doi:10.1038/tp.2014.120 (2014).

32 McNamara, I. M., Borella, A. W., Bialowas, L. A. & Whitaker-Azmitia, P. M. Further studies in the developmental hyperserotonemia model (DHS) of autism: social, behavioral and peptide changes. *Brain Res* **1189**, 203-214, doi:10.1016/j.brainres.2007.10.063 (2008).

56 Launay, J. M. *et al.* One-step purification of the serotonin transporter located at the human platelet plasma membrane. *J Biol Chem* **267**, 11344-11351 (1992).

57 Launay, J. M., Geoffroy, C., Costa, J. L. & Alouf, J. E. Purified -SH-activated toxins (streptolysin O, alveolysin): new tools for determination of platelet enzyme activities. *Thromb Res* **33**, 189-196, doi:10.1016/0049-3848(84)90179-8 (1984).

58 Launay, J. M. & Alouf, J. E. Biochemical and ultrastructural study of the disruption of blood platelets by streptolysin O. *Biochim Biophys Acta* **556**, 278-291, doi:10.1016/0005-2736(79)90048-8 (1979).

59 Fuertig, R. *et al.* LC-MS/MS-based quantification of kynurenine metabolites, tryptophan, monoamines and neopterin in plasma, cerebrospinal fluid and brain. *Bioanalysis* **8**, 1903-1917, doi:10.4155/bio-2016-0111 (2016).

60 Trammell, S. A. *et al.* Nicotinamide riboside is uniquely and orally bioavailable in mice and humans. *Nat Commun* **7**, 12948, doi:10.1038/ncomms12948 (2016).

61 Namboodiri, M. A., Nakai, C. & Klein, D. C. Effects of selected treatments on stability and activity of pineal serotonin N-acetyltransferase. *J Neurochem* **33**, 807-810, doi:10.1111/j.1471-4159.1979.tb05229.x (1979).

62 Axelrod, J., Wurtman, R. J. & Snyder, S. H. Control of Hydroxyindole O-Methyltransferase Activity in the Rat Pineal Gland by Environmental Lighting. *J Biol Chem* **240**, 949-954 (1965).

63 Ben-Abdallah, M. *et al.* Production of soluble, active acetyl serotonin methyl transferase in Leishmania tarentolae. *Protein Expr Purif* **75**, 114-118, doi:10.1016/j.pep.2010.07.011 (2011).

64 Ferry, G. *et al.* Purification of the recombinant human serotonin N-acetyltransferase (EC 2.3.1.87): further characterization of and comparison with AANAT from other species. *Protein Expr Purif* **38**, 84-98, doi:10.1016/j.pep.2004.07.004 (2004).

65 Tugaeva, K. V., Remeeva, A., Gushchin, I., Cooley, R. B. & Sluchanko, N. N. Design, expression, purification and crystallization of human 14-3-3zeta protein chimera with phosphopeptide from proapoptotic protein BAD. *Protein Expr Purif* **175**, 105707, doi:10.1016/j.pep.2020.105707 (2020).

66 Walker, R. F., Friedman, D. W. & Jimenez, A. A modified enzymatic-isotopic microassay for serotonin (5HT) using 5HT-N-acetyltransferase partially purified from Drosophila. *Life Sci* **33**, 1915-1924, doi:10.1016/0024-3205(83)90676-8 (1983).

67 Nishikawa, M. *et al.* Whole-cell-dependent biosynthesis of sulfo-conjugate using human sulfotransferase expressing budding yeast. *Appl Microbiol Biotechnol* **102**, 723-732, doi:10.1007/s00253-017-8621-x (2018).

68 Falany, C. N. & Kerl, E. A. Sulfation of minoxidil by human liver phenol sulfotransferase. *Biochem Pharmacol* **40**, 1027-1032, doi:10.1016/0006-2952(90)90489-8 (1990).

69 Malherbe, P. *et al.* Molecular cloning and functional expression of human 3-hydroxyanthranilic-acid dioxygenase. *J Biol Chem* **269**, 13792-13797 (1994).

70 Vittet, D., Rondot, A., Cantau, B., Launay, J. M. & Chevillard, C. Nature and properties of human platelet vasopressin receptors. *Biochem J* **233**, 631-636, doi:10.1042/bj2330631 (1986).

71 Gould, G. G., Barba-Escobedo, P. A., Horton, R. E. & Daws, L. C. High Affinity Decynium-22 Binding to Brain Membrane Homogenates and Reduced Dorsal Camouflaging after Acute Exposure to it in Zebrafish. *Front Pharmacol* **13**, 841423, doi:10.3389/fphar.2022.841423 (2022).

72 Delorme, R. *et al.* Platelet serotonergic markers as endophenotypes for obsessive-compulsive disorder. *Neuropsychopharmacology* **30**, 1539-1547, doi:10.1038/sj.npp.1300752 (2005).

73 Cesura, A. M., Ritter, A., Picotti, G. B. & Da Prada, M. Uptake, release, and subcellular localization of 1-methyl-4-phenylpyridinium in blood platelets. *J Neurochem* **49**, 138-145, doi:10.1111/j.1471-4159.1987.tb03405.x (1987).

74 Mony, T. J., Lee, J. W., Dreyfus, C., DiCicco-Bloom, E. & Lee, H. J. Valproic Acid Exposure during Early Postnatal Gliogenesis Leads to Autistic-like Behaviors in Rats. *Clin Psychopharmacol Neurosci* **14**, 338-344, doi:10.9758/cpn.2016.14.4.338 (2016).

75 Mirza, R. & Sharma, B. Benefits of Fenofibrate in prenatal valproic acid-induced autism spectrum disorder related phenotype in rats. *Brain Res Bull* **147**, 36-46, doi:10.1016/j.brainresbull.2019.02.003 (2019).

76 Kumar, H. & Sharma, B. Memantine ameliorates autistic behavior, biochemistry & blood brain barrier impairments in rats. *Brain Res Bull* **124**, 27-39, doi:10.1016/j.brainresbull.2016.03.013 (2016).

77 Mony, T. J., Hong, M. & Lee, H. J. Empathy Study in Rodent Model of Autism Spectrum Disorders. *Psychiatry Investig* **15**, 104-110, doi:10.30773/pi.2017.06.20 (2018).

78 Madden, A. M. & Zup, S. L. Effects of developmental hyperserotonemia on juvenile play behavior, oxytocin and serotonin receptor expression in the hypothalamus are age and sex dependent. *Physiol Behav* **128**, 260-269, doi:10.1016/j.physbeh.2014.01.036 (2014).

79 Lai, M. C., Lombardo, M. V. & Baron-Cohen, S. Autism. *Lancet* **383**, 896-910, doi:10.1016/S0140-6736(13)61539-1 (2014).

80 Mirza, R. & Sharma, B. Selective modulator of peroxisome proliferator-activated receptor-alpha protects propionic acid induced autism-like phenotypes in rats. *Life Sci* **214**, 106-117, doi:10.1016/j.lfs.2018.10.045 (2018).

81 Veenstra-VanderWeele, J. *et al.* Autism gene variant causes hyperserotonemia, serotonin receptor hypersensitivity, social impairment and repetitive behavior. *Proc Natl Acad Sci U S A* **109**, 5469-5474, doi:10.1073/pnas.1112345109 (2012).

82 Cuevas-Olguin, R. *et al.* Cerebrolysin prevents deficits in social behavior, repetitive conduct, and synaptic inhibition in a rat model of autism. *J Neurosci Res* **95**, 2456-2468, doi:10.1002/jnr.24072 (2017).

### Legends to the Supplementary Figures

**Extended Fig. 1: TMAO in mice administered with exogenous TMAO or FMO3 inhibitor. a,** plasma (**a**) and platelet (**b**) TMAO levels in wild-type mice (n = 5) at baseline and after the successive administration of exogenous TMAO and FMO3 inhibitor. Intergroup comparisons were performed using the Kruskal-Wallis test followed by the Wilcoxon sum-rank test corrected for multiple comparisons.

**Extended Fig. 2: V1aR concentration in the platelets** of controls (n = 106) and individuals with ASD (n = 90). Intergroup comparisons were performed using the Wilcoxon sum-rank test.

**Extended Fig. 3: Evolution of biochemical and behavioural parameters after withdrawal of urea in DHS rats. a-e,** Plasma sodium (**a**), plasma TMAO (**b**), whole-blood serotonin (**c**), plasma melatonins (**d**), and plasma $NAD^+$ (**e**) concentrations in DHS rats that received urea from PDN21-25 (n = 8) at PDN33 (D33) , PDN60 (D60), and PDN90 (D90). **f-l,** sociability index (**f**), social preference index (**g**), time spent in the open arm (TSOA, **h**) and the number of open arm entries (OAE, **i**) of the elevated plus maze (EPM), number of line crossing (**j**) and central square entries (**k**) of the open field (OF), and spontaneous alternation (**l**) in DHS rats that received urea from PDN21-25 at PDN33 (D33) , PDN60 (D60), and PDN90 (D90, n = 8). Within-group comparisons were performed using repeated-measure ANOVA on log-transformed data followed by the Tukey HSD test. Of note, the values at PDN33 are those depicted in **Fig. 4 b-l** as DHS + urea, as the same animals were used.

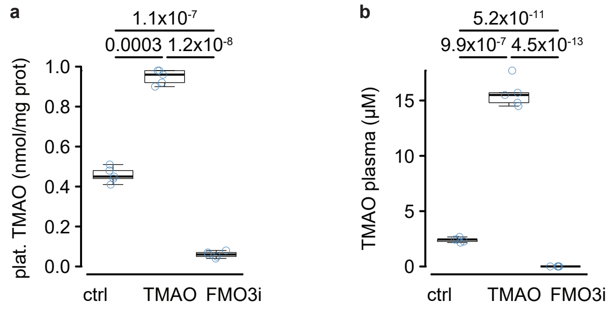


**Extended Fig. 1: TMAO in mice administered with exogenous TMAO or FMO3 inhibitor. a,** plasma (**a**) and platelet (**b**) TMAO levels in wild-type mice (n = 5) at baseline and after the successive administration of exogenous TMAO and FMO3 inhibitor. Intergroup comparisons were performed using the Kruskal-Wallis test followed by the Wilcoxon sum-rank test corrected for multiple comparisons.

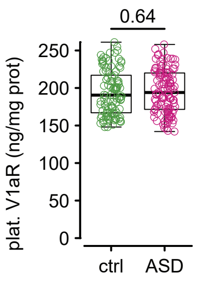


**Extended Fig. 2:** V1aR concentration in the platelets of controls (n = 106) and individuals with ASD (n = 90). Intergroup comparisons were performed using the Wilcoxon sum-rank test.

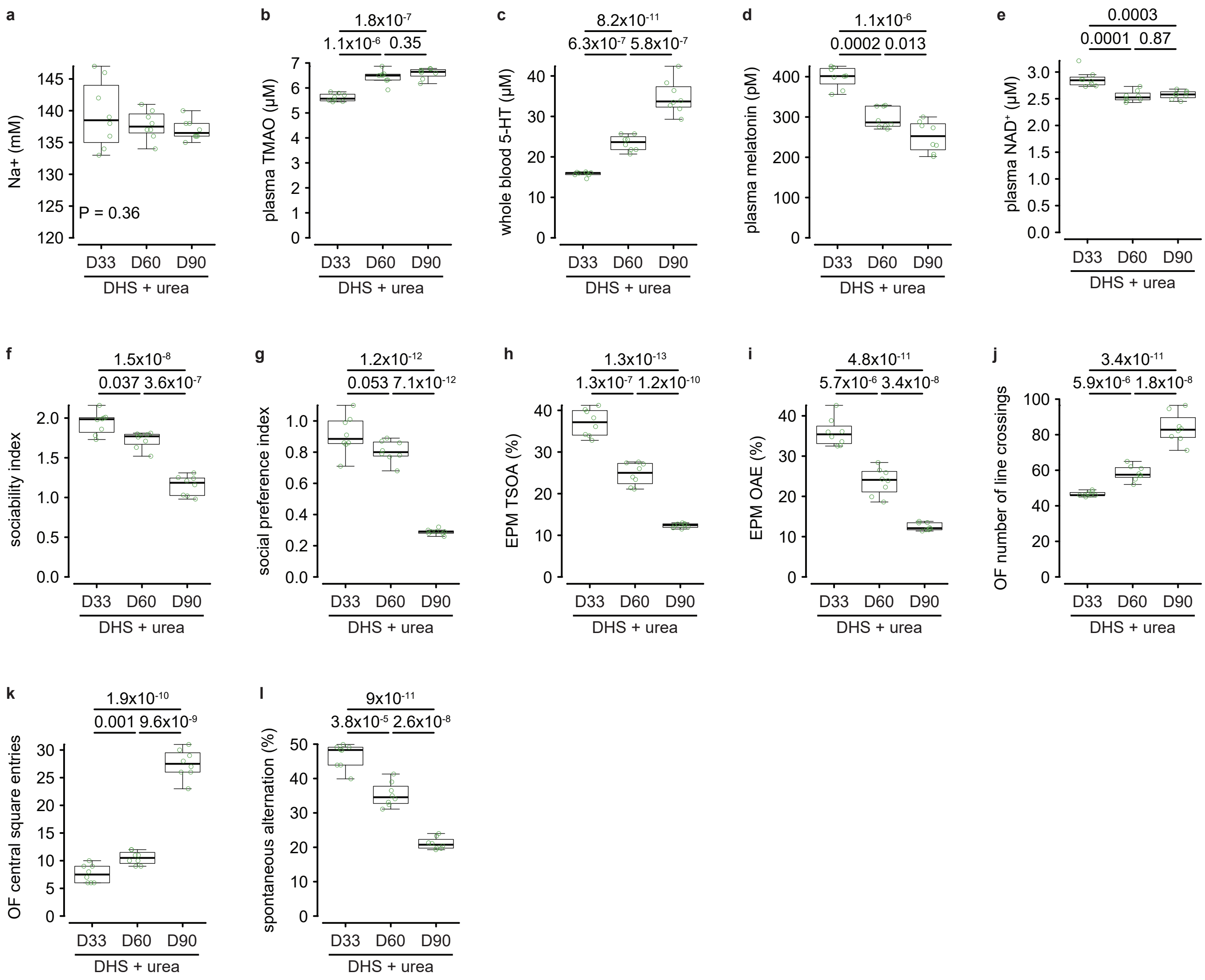


**Extended Fig. 3: Evolution of biochemical and behavioural parameters after withdrawal of urea in DHS rats. a-e,** Plasma sodium (**a**), plasma TMAO (**b**), whole-blood serotonin (**c**), plasma melatonins (**d**), and plasma $NAD^+$ (**e**) concentrations in DHS rats that received urea from PDN21-25 (n = 8) at PDN33 (D33) , PDN60 (D60), and PDN90 (D90). **f-l,** sociability index (**f**), social preference index (**g**), time spent in the open arm (TSOA, **h**) and the number of open arm entries (OAE, **i**) of the elevated plus maze (EPM), number of line crossing (**j**) and central square entries (**k**) of the open field (OF), and spontaneous alternation (**l**) in DHS rats that received urea from PDN21-25 at PDN33 (D33) , PDN60 (D60), and PDN90 (D90, n = 8). Within-group comparisons were performed using repeated-measure ANOVA on log-transformed data followed by the Tukey HSD test. Of note, the values at PDN33 are those depicted in **Fig. 4 b-l** as DHS + urea, as the same animals were used.
